# dearseq: a variance component score test for RNA-Seq differential analysis that effectively controls the false discovery rate

**DOI:** 10.1101/635714

**Authors:** Marine Gauthier, Denis Agniel, Rodolphe Thiébaut, Boris P. Hejblum

## Abstract

RNA-seq studies are growing in size and popularity. We provide evidence that the most commonly used methods for differential expression analysis (DEA) may yield too many false positive results in some situations. We present `dearseq`, a new method for DEA which controls the FDR without making any assumption about the true distribution of RNA-seq data. We show that `dearseq` controls the FDR while maintaining strong statistical power compared to the most popular methods. We demonstrate this behavior with mathematical proofs, simulations, and a real data set from a study of Tuberculosis, where our method produces fewer apparent false positives.

## Background

With the rise of next generation sequencing technologies that measure gene expression on the scale of the entire genome, RNA-seq differential expression analysis (DEA) has become ubiquitous in many research fields. While numerous approaches have been proposed to perform DEA of RNA-seq data, there is no clear consensus on which method is the most efficient. Three methods stand out as the most commonly used in practice: `edgeR` [1], `DESeq2` [2], and `limma-voom` [3] (respectively 5,853, 5,019, and 812 citations in PubMed as of May 15^th^, 2019). `edgeR` and `DESeq2` both rely on the assumption that gene counts from RNA-seq measurements follow a negative binomial distribution, and `limma-voom` is based on a weighted linear model and assumes resulting test statistics follow a normal distribution.

Following long-standing statistical practice, researchers typically attempt to control the probability of finding a gene to be differentially expressed (DE) when the opposite is true in reality (i.e. the Type-I error) at a pre-specified level (conventionally 5%). In a high-dimensional context such as gene expression data, the false positive rate or False Discovery Rate (FDR) [4] has been largely adopted as the target probability to be controlled in exploratory studies. The FDR is the expected proportion of features identified as significant that are actually false positive: for instance, an FDR of 5% implies that among all the genes declared DE, 5% are not DE. Controlling this error rate results in many fewer false positives than controlling the per-gene Type-I error, while not being as restrictive as controlling the probability of any false positive (the family-wise error rate) among all of the potentially thousands of genes.

This control is usually taken for granted and often left out from the benchmarks of DEA methods, while in fact, an excessive FDR can be quite problematic. Not controlling the FDR means getting more false positives than expected, which limits the reproducibility of study results. Whole genome DEAs are usually exploratory steps prior to subsequent studies to confirm a gene signature is associated with a particular biological condition. If a majority of the selected genes turn out to be false positives, results may fail to replicate and any down-stream health benefits may remain elusive, not to mention the waste of precious research resources.

When comparing DEA methods, the evaluation of their empirical FDR with respect to the targeted (nominal) level is often overlooked[5, 6, 7, 8, 9, 10]. Nonetheless, some issues with inflated FDR in DEA have been previously reported in the literature [11, 12, 13, 14, 15], but those warnings have made little apparent impact on DEA practices.

Inflation of the empirical FDR in DEA can have numerous causes, from inadequate preprocessing of the data to violations of the DEA method’s underlying assumptions. In particular, `edgeR`, `DESeq2` and `limma-voom` make potentially strong distributional assumptions on RNA-seq data. This type of model-based inference may be required when RNA-seq studies include only a small number of observations. However, these methods’ parametric assumptions are not typically verifiable in practice. Any deviation from the hypothesized distribution of test statistics will translate into ill-behaved *p*-values and therefore uncontrolled FDR. FDR control rests upon the entire distribution of *p*-values being uniform under the null hypothesis *H*_0_ (i.e. for genes that are truly not DE). So even a slight deviation from strict Type-I error control can have dramatic consequences on the empirical FDR. In addition, even if Type-I error were controlled at say 5%, non-uniformity in the *p*-value distribution could lead to failure to control the Type-I error at lower levels (such as 1% or lower) and/or failure to control the FDR. When problems with *p*-values and FDR arise due to violation of modeling assumptions, larger sample sizes will only exacerbate the problem. As sequencing costs keep falling, study sample sizes are increasing, making this issue more urgent.

Here, we propose `dearseq`, a new method to perform DEA that effectively controls the FDR, regardless of the distribution of the underlying data. `dearseq` is a robust approach that uses a variance component score test and relies on nonparametric regression to account for the intrinsic heteroscedasticity of RNAseq data. In the Results section we compare the performance of `dearseq` to the three most popular state-of-the-art methods for DEA: `edgeR`, `DESeq2` and `limma-voom`. We demonstrate that `dearseq` enforces strict control of Type-I error and FDR while maintaining good statistical power in a realistic and extensive simulation study where knowing the ground truth facilitates benchmarking the properties of the different methods. We also present a comparative re-analysis of a real-world Tuberculosis data set from Singhania *et al.* [16] studying apparent false positives identified by the leading DEA methods compared to `dearseq`. `dearseq` can efficiently identify the genes whose expression is significantly associated with one or several factors of interest in complex experimental designs (including longitudinal observations) from RNA-seq data while providing robust control of FDR. `dearseq` is freely available as an R package on the Bioconductor library.

## Results

### Synthetic simulation study

As highlighted by both Conesa *et al.*[17] and Assefa *et al.* [15], engaging in realistic yet clear simulations is difficult. One has to find the right balance between the controlled settings necessary to know the ground truth, and the realism necessary to be convincing that the results would translate in real-world analyses. In an attempt to cover as broad a spectrum as possible, we present a performance evaluation of our methods under three data-generating scenarios: a) a negative binomial parametric assumption for RNA-seq data, b) a highly non-linear model designed to violate most modeling assumptions, and c) a resampling from SEQC data [18] with truncated Gaussian noise. Scenario a) may be favorable to `edgeR` and `DESeq2` as it relies on their parametric assumption of a negative binomial distribution for RNA-seq count data. Scenario b) may be unfavorable for all three compared methods (`edgeR`, `DESeq2` and `limma-voom`) since it features a high degree of non-linearity, deviating from any assumed model. Scenario c) is likely the most realistic of the three because it relies only on resampling real RNA-seq samples from the SEQC study[18], similarly to what was done in Germain *et al.* [13]. A multivariate truncated Gaussian noise (using the estimated covariance stucture across the observed genes) was added to enable the generation of larger sample sizes while preserving the count nature of the data.

We simulated 1,000 synthetic data sets at different sample sizes using each one of these three scenarios. For scenarios a) and b), 10% of genes were generated as truly DE while the remaining 90% were not DE. For the scenario c), since it is based on resampling from homogeneous samples, it was impossible to induce truly DE genes without making further parametric assumptions (which would have made the scenario less realistic). For this reason, in scenario c), FDR corresponded to the probability of finding any genes to be DE. Details of the data-generating mechanisms are provided in Additional file 1.

We evaluated the four methods (the leading methods and `dearseq`) in terms of Type-I error control and statistical power, as well as in terms of FDR and True Discovery Rate (TDR) after Benjamini-Hochberg [4] correction for multiple testing. Throughout, we used a targeted control rate for the FDR at a nominal level of 5%. Fig. 1 presents the Monte-Carlo estimation over the 1,000 simulations in each of the three scenarios for both the Type-I error and the FDR according to increasing samples sizes (from 4 to 300 samples). Fig. 2 presents the results of the first two scenarios for both the statistical power and the TDR.

**Figure 1:**
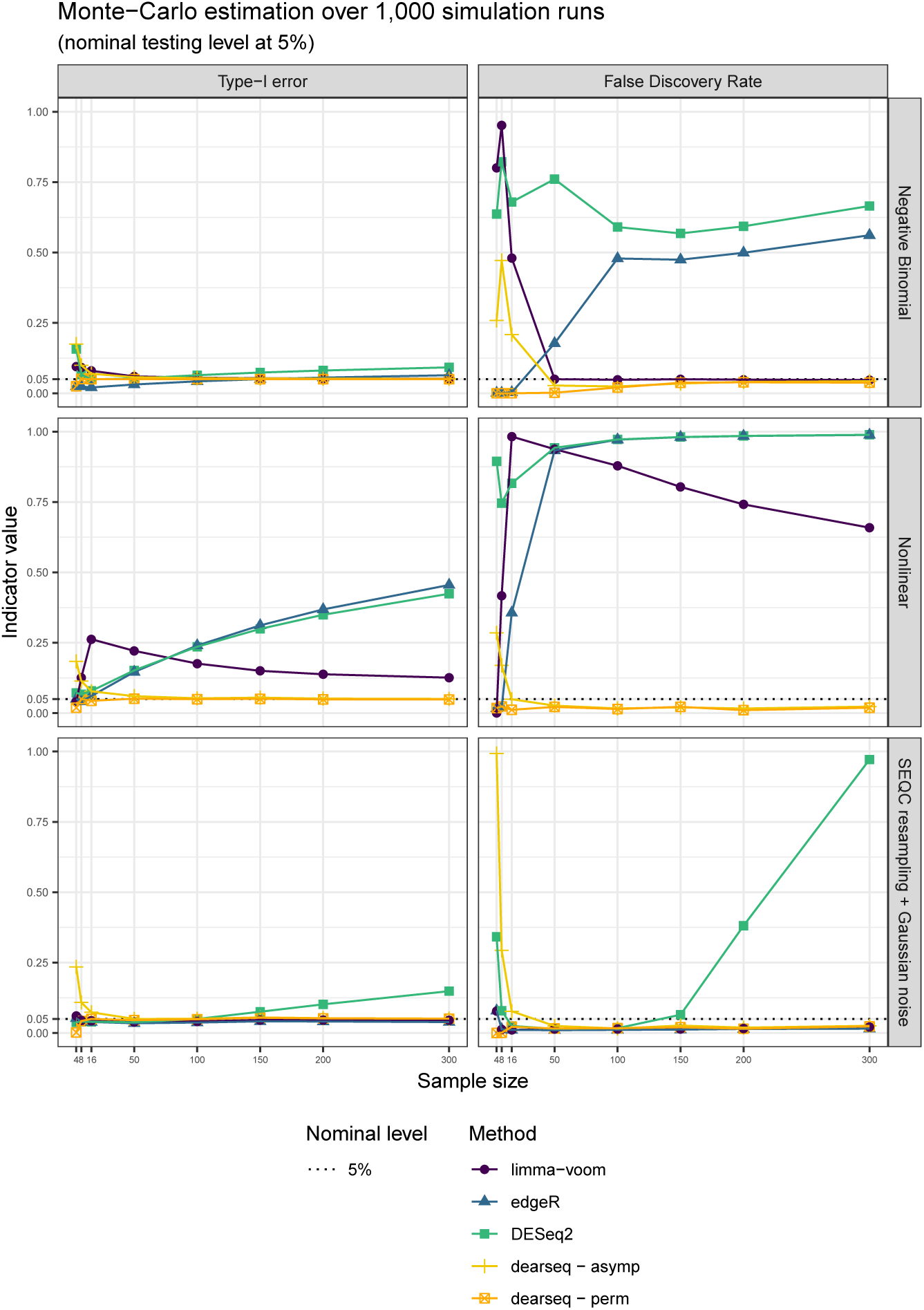
Type-I error and FDR curves for each DEA method with increasing sample sizes. In each setting (Negative Binomial, Non-linear, and SEQC data resampling), the Type-I error is computed as the number significant genes among the true negative, and the FDR as the average number of false positives among the genes declared DE.

**Figure 2:**
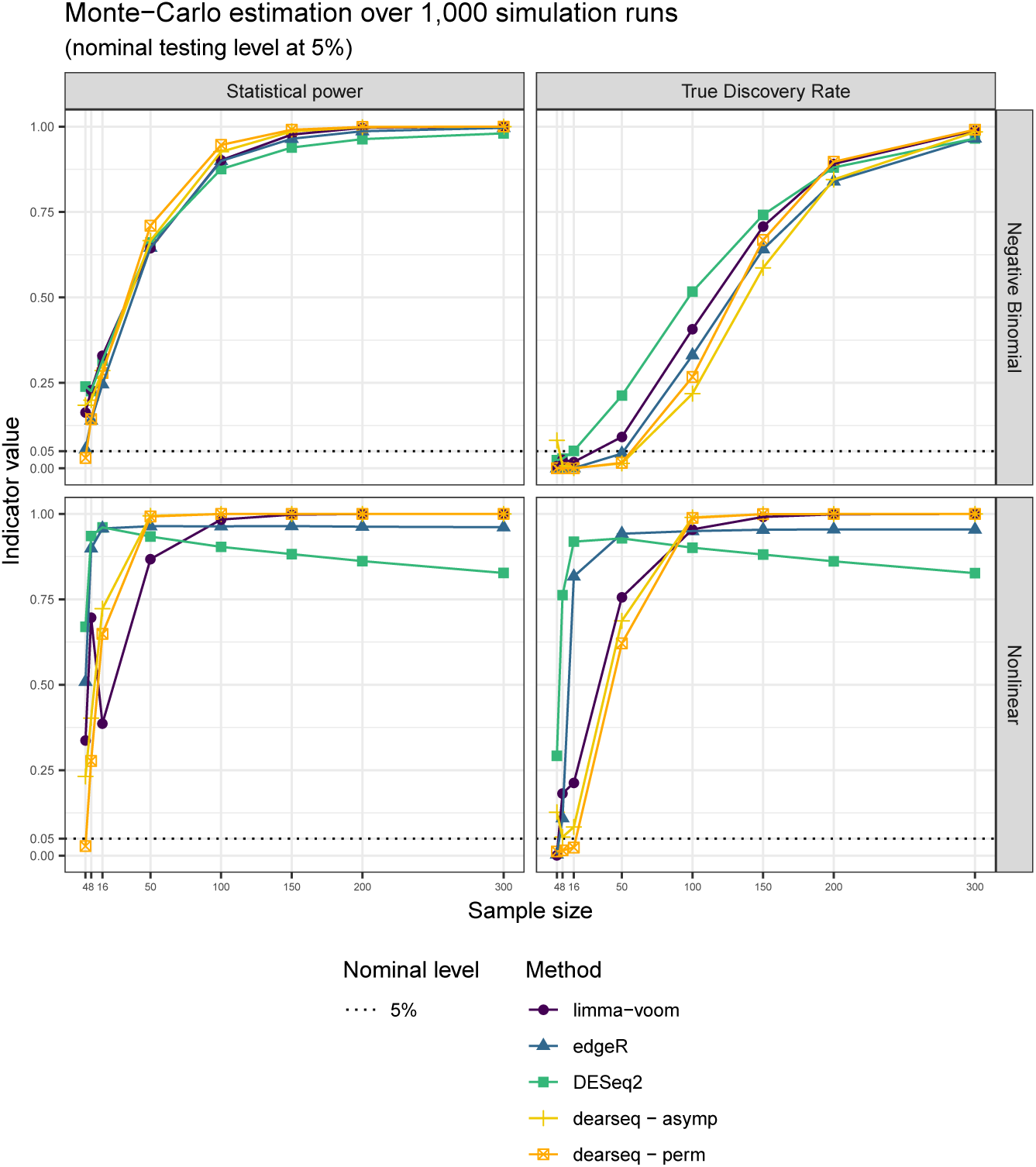
Power and True Discovery Rate curves for each DEA method with increasing sample size. Because SEQC data resampling only generates non-significant genes, this setting does not allow to estimate statistical power or TDR.

In Fig. 1, `dearseq` exhibited good control of both Type-I error and FDR in all three scenarios, as soon as asymptotic convergence was reached (between 16 and 50 samples depending on the scenarios). To accommodate small sample sizes, we have also developed a permutation-based version of `dearseq`, which always controlled Type-I error and FDR, regardless of the sample size. `edgeR` appeared to control the Type-I error in both scenarios a) and c), but exhibits slightly inflated Type-I error for large sample sizes in scenario a) (from 100 samples). This was much more visible for the FDR, which `edgeR` failed to control as soon as the sample size rose above 50. Under the non-linear model, neither the Type-I error nor the FDR are controlled by `edgeR`. `limma-voom` exhibited good control of both the Type-I error and FDR as long as its linear hypothesis was not violated (i.e., not scenario b)) and the sample size was large enough (between 8 to 50 samples depending on the scenario). Finally, `DESeq2` failed to control either the Type-I error or the FDR in any of the three scenarios, with its problems growing worse as the sample size increased.

Fig. 2 shows that this robust control of Type-I error and FDR from `dearseq` does not come at a price of reduced statistical power (or True Discovery Rate, its multiple-testing correction equivalent). Interestingly, the permutation approach also exhibit good statistical power. Regarding competing approaches, interpreting statistical power when the Type-I error is not controlled would be a bit dubious.

### Real data set

In a recent paper, Singhania *et al.* identified a 373-genes signature of active tuberculosis from RNA-seq data [16]. Tuberculosis (TB) is a disease caused by a bacterium called Mycobacterium tuberculosis. Bacteria typically infect the lungs, but they can also affect other parts of the body. Tuberculosis can remain in a quiescent state called latent tuberculosis infection (LTBI), where the patient is infected but has no clinical, bacteriological or radiological signs of the disease. Participants to this study were recruited from several medical institutes in London, UK (see Berry *et al.* [19] for a detailed description). All participants were aged over 18 years old. Active TB patients were confirmed by laboratory isolation of M. tuberculosis on mycobacterial culture of a respiratory specimen, while Latent TB patients were characterized by a positive tuberculin-skin test (TST) together with a positive result using a M. tuberculosis antigen specific IFN-*γ* release assay (IGRA). Healthy control participants were recruited from volunteers at the National Institute for Medical Research (NIMR, Mill Hill, London, UK) and were negative to both TST and IGRA. In total, 54 participants were included, of whom 21 were active TB patients, 21 were LTBI patients, and 12 were healthy controls.

The signature was derived by contrasting active tuberculosis (TB) patients on the one hand against healthy individuals (Control) or those with a latent infection (LTBI) on the other hand (see Fig. 3). Their original analysis applied `edgeR` to their Berry London RNA-seq data, which included 14,150 normalized-gene counts measured across 54 samples after preprocessing (see Singhania *et al.* or supplementary material in additional file 1 for more information on this preprocessing) available from GEO (GSE107991). In light of our simulation results regarding false positives, we sought to investigate how many of the 373 genes Singhania *et al.* found using `edgeR` might actually be false positives. We therefore conducted a comparative re-analysis of these data, first comparing DE genes found by `dearseq` to the original signature of Singhania *et al.*. Secondly, we further compared the results obtained from the other leading methods, `DESeq2` and `limma-voom`.

**Figure 3:**
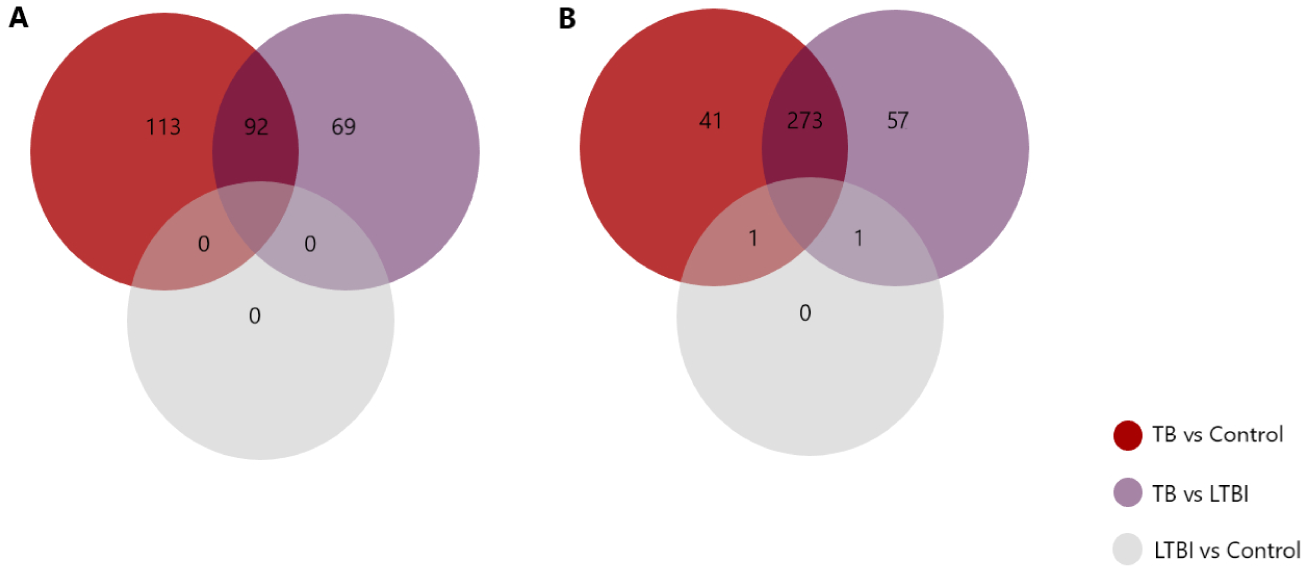
Venn diagram showing overlap of DE genes using dearseq and the original edgeR signature among the three comparisons performed. a) Venn diagram showing the results of the three DEA using `dearseq`. Note that no gene differentially expressed was found with our method comparing the LTBI group and the control group, unlike `edgeR` which found two such genes to be DE. b) Venn diagram showing the results of the DEA using `edgeR` (Singhania *et al.*).

Following Singhania *et al.*, to be included in the signature a DE gene *g* must have had both: i) an absolute log`2`(fold change) *>* 1, and ii) an FDR adjusted *p*-value *<* 0.05 (after correction for multiple testing with the Benjamini-Hochberg procedure). To ensure reproducibility of the numerical values from Singhania *et al.*, the log`2` fold changes were calculated using `edgeR`. The signature was then evaluated by its capacity to distinguish between active TB versus all others. In order to quantify the relevance of each selected gene for distinguishing active TB from control and LTBI, we computed two measures of association, the leave-one-out cross-validated Brier score [20] and the marginal *p*-value for the association between the gene and TB status. The Brier score was computed as 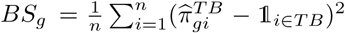. It compares each patient’s TB status to 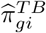;, their predicted probability of TB based on the selected gene *g* estimated using leave-one-out cross validation. A gene with Brier score *BS*_*g*_ close to 0 is a good predictor of TB, while a gene with Brier score far away from 0 is a poor predictor and potential false positive. Similarly, we compute the marginal *p*-value for each gene from a logistic regression predicting TB status from the gene expression. We estimate the Brier score and *p*-value for each gene separately. We do this rather than a multivariate model including all genes because the presence of a single predictive gene in the multivariate signature would be enough to yield accurate predictions, thus masking the potential false positive genes included in the model.

Applying the `dearseq` permutation test (see Methods) to the three comparisons originally performed in Singhania *et al.* (TB vs Control, LTBI vs Control, and TB vs LTBI) yields a global signature of 274 DE genes (see Fig. 3) of which 234 are in common with those found by the original `edgeR` analysis (see Fig. 5). We isolated the genes only identified by `dearseq` from the genes only identified by `edgeR` and from the genes in common between the two signatures to further assess the differences between the two results. Comparing the gene specific Brier scores *BS*_*g*_ between the two signatures clearly shows that the overwhelming majority of the highest scores (i.e. the lowest predictive abilities) is due to `edgeR`-private genes (see Fig. 4 b)). Indeed, the univariate Brier scores of the `dearseq`-private genes have significatively smaller values on average than the `edgeR`-private genes (according to a *t*-test – see Fig. 4). This is further confirmed by the marginal association *p*-values, for which all of the highest values are again from `edgeR` private, notably all the values above 0.05. Thus, `edgeR`-private genes are likely false positives whereas the `dearseq`-private sound more relevant. In a biological point of view, the main pathways concerned by the 139 `edgeR`-private genes, that are “Inhibition of matrix metalloproteinases”, “Granulocyte Adhesion and Diapedesis”, “Inhibition of Angiogenesis by TSP1” using Ingenuity Pathways Analysis (IPA) were not directly related to the main pathways observed in the retained 373-gene signature (IFN-inducible genes, B- and T-cell genes). Those results emphasize the better predictive ability of the genes identified by `dearseq`, and highlights the potential false positives arising from `edgeR`.

**Figure 4:**
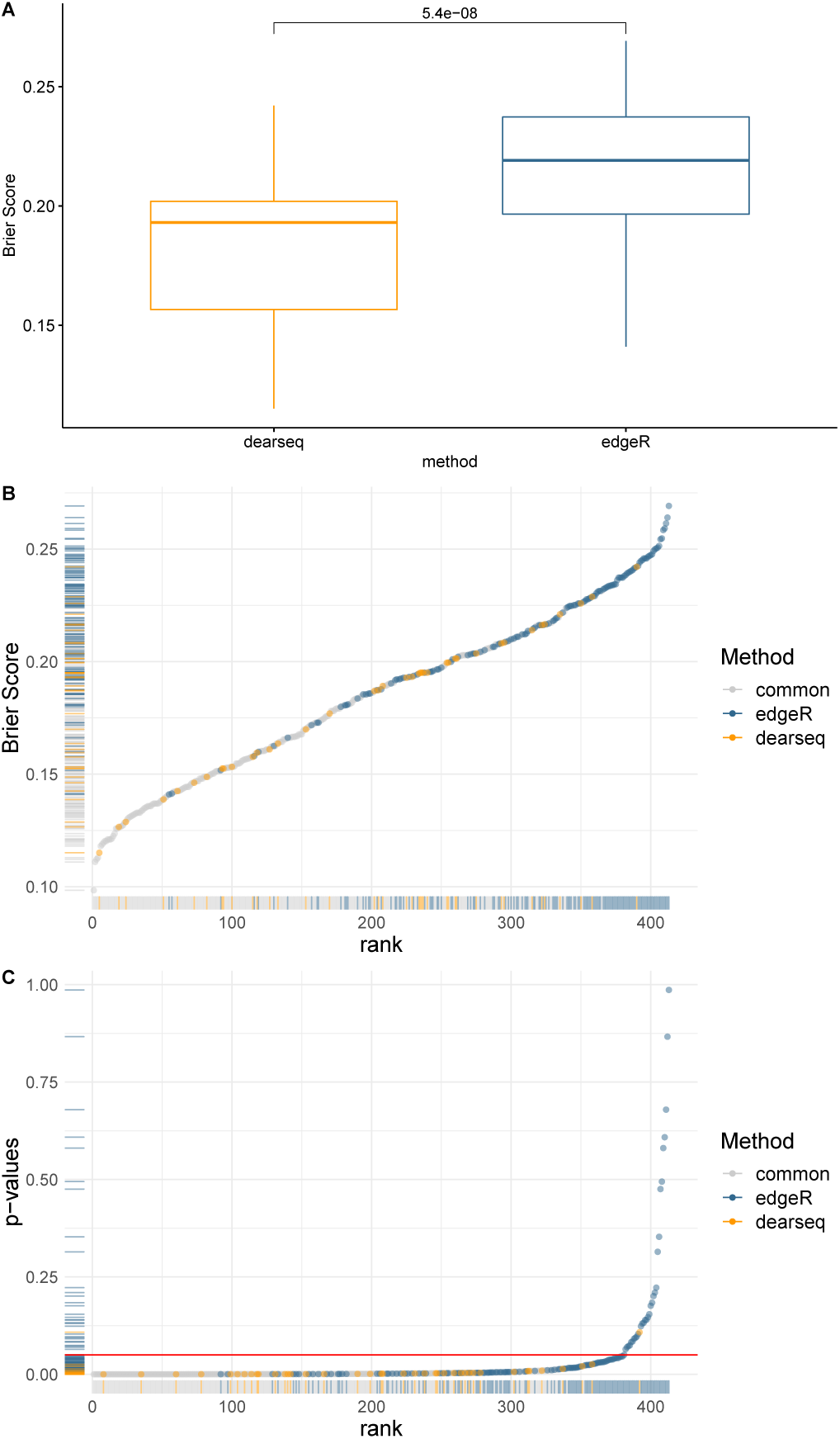
Comparing edgeR-based signature to the signature derived by dearseq. a) Boxplots of the Brier scores of the 40 genes private to `dearseq` (i.e., not also declared DE by `edger`) and the 139 genes private to the original `edgeR` analysis. b) Univariate Brier scores. The blue points correspond to genes found only in the original `edgeR` signature, the yellow points found only in the `dearseq` signature, and the grey points found in both signatures. c) Marginal *p*- values from a univariate logistic regression combined with a leave-one-out cross validation for the 40 `dearseq`-private and the 139 `edgeR`-private genes. The red line indicates the common 5% *p*-value threshold.

**Figure 5:**
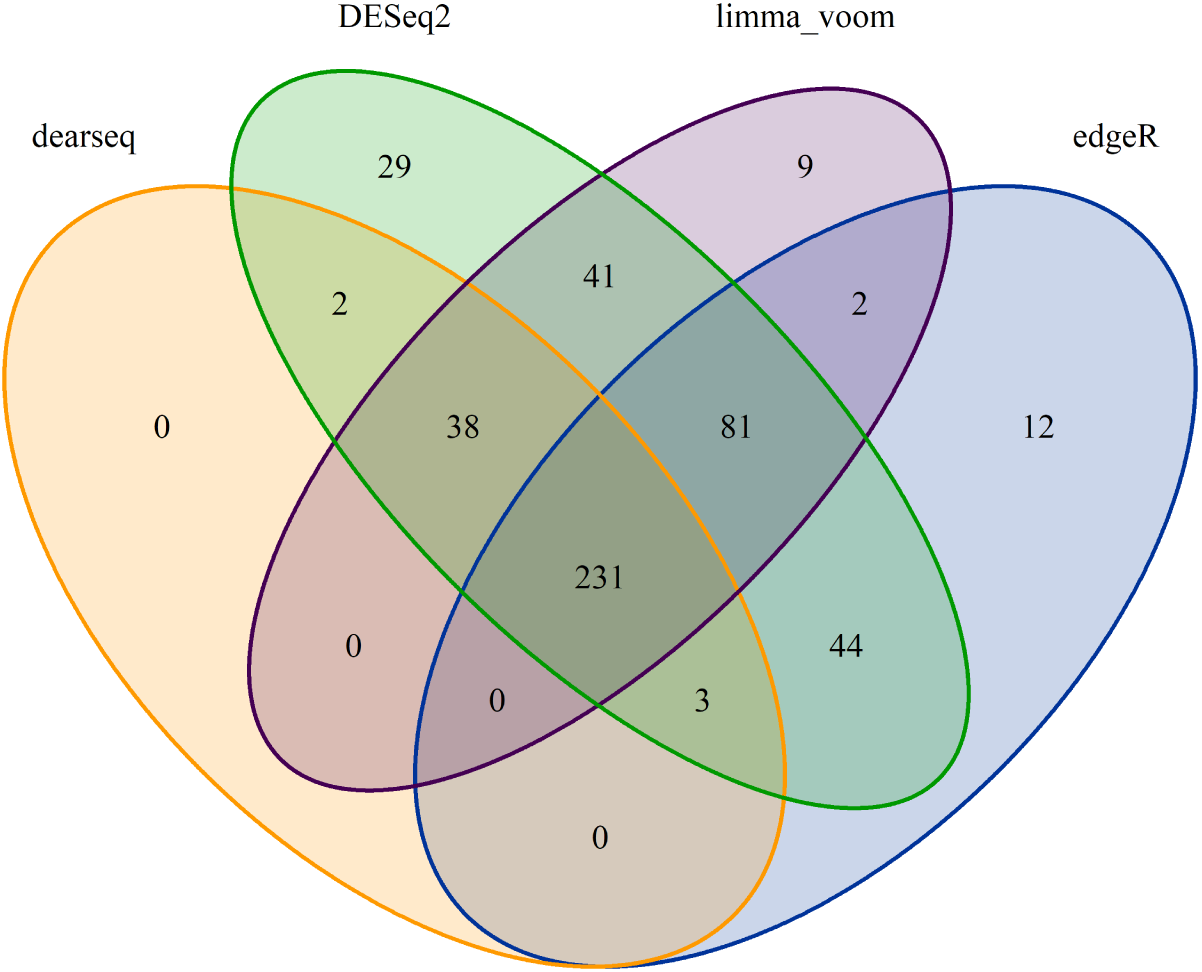
Venn diagram summarizing the different signatures from the four methods. Venn diagram of the genes declared DE by `dearseq`, `DESeq2`, `limma-voom` and `edgeR` (Singhania *et al.*) under an FDR-adjusted *p*-value of 0.05. None of the genes is found with `dearseq` only.

In addition, we also performed the same analysis using `limma-voom` and `DESeq2` to further benchmark the performance of `dearseq`. Fig. 5 displays the Venn diagram of significantly DE genes across these four analyses. There are 231 genes common across all these tools. Interestingly, all of the 274 genes identified by `dearseq` are also identified by at least one of the three competing methods (and only 2 genes are idendified by less than two other methods – namely only by `DESeq2`), illustrating that `dearseq` is less prone to generate false positives. `DESeq2` identifies the largest signature comprising of 457 genes, including all of the 274 genes identified by `dearseq` and 359 out of the 373 originally identified by `edgeR`, while `limma-voom` identify 402 genes, among which 269 are in common with `dearseq` and 314 are in common with `edgeR`. As can be seen on Fig. 6 the `dearseq` signature has the lowest average Brier score, meaning that most of the additional genes identified by the three competing methods are less predictive of active TB status. Fig. 7 strengthens this conclusion by showing again that the `limma-voom`-private are largely over-represented among the highest Brier scores and the highest marginal *p*-values. The same conclusion can be drawn for the `DESeq2`-private genes.

**Figure 6:**
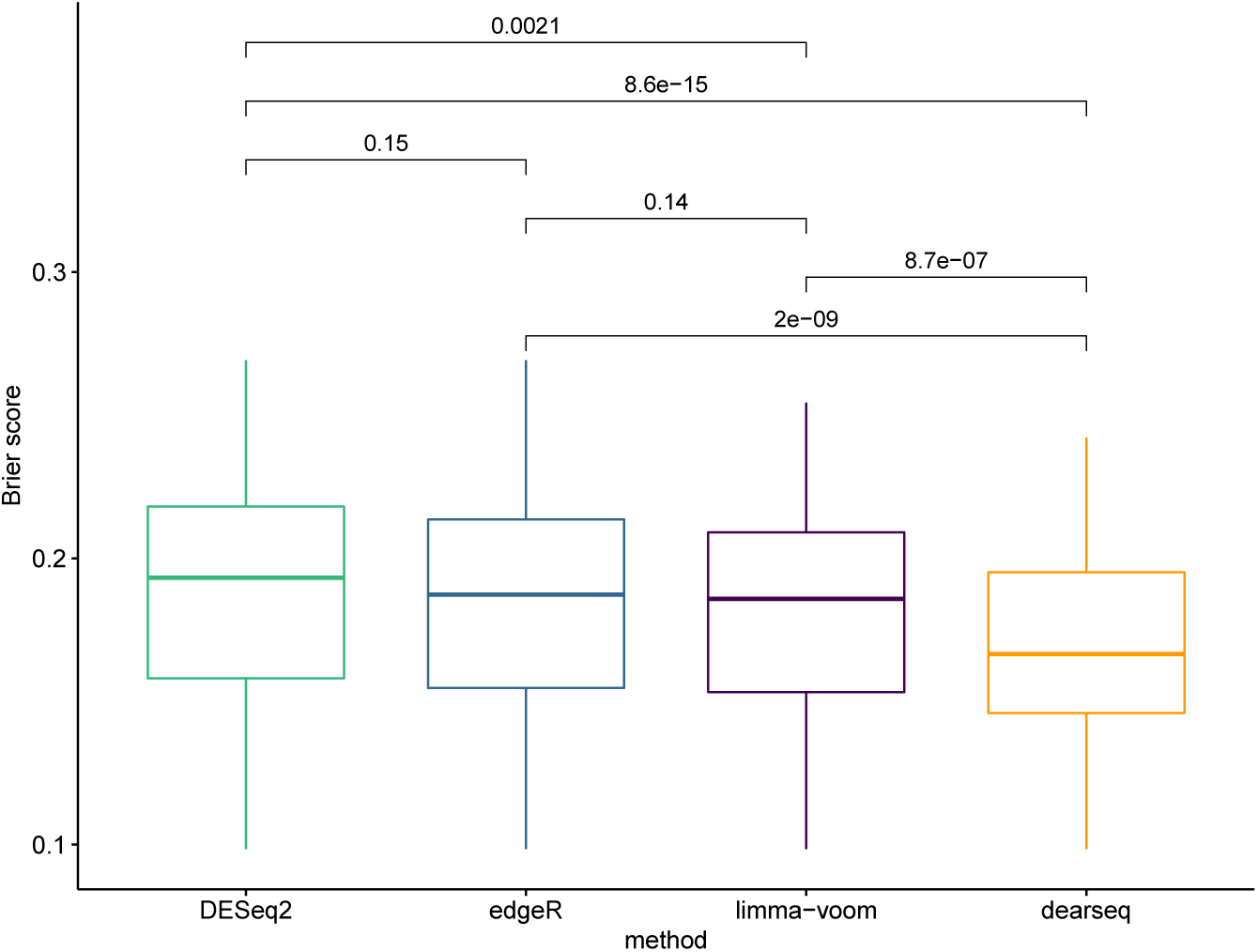
Boxplots of the Brier scores of all the genes declared DE by the four methods. Boxplots of the Brier scores of all the DE genes called by `dearseq`, `DESeq2`, `limma-voom` and `edgeR` (Singhania *et al.*). The predictions are derived from a logistic regression combined with a leave-one-out cross validation. Smaller Brier scores are better.

**Figure 7:**
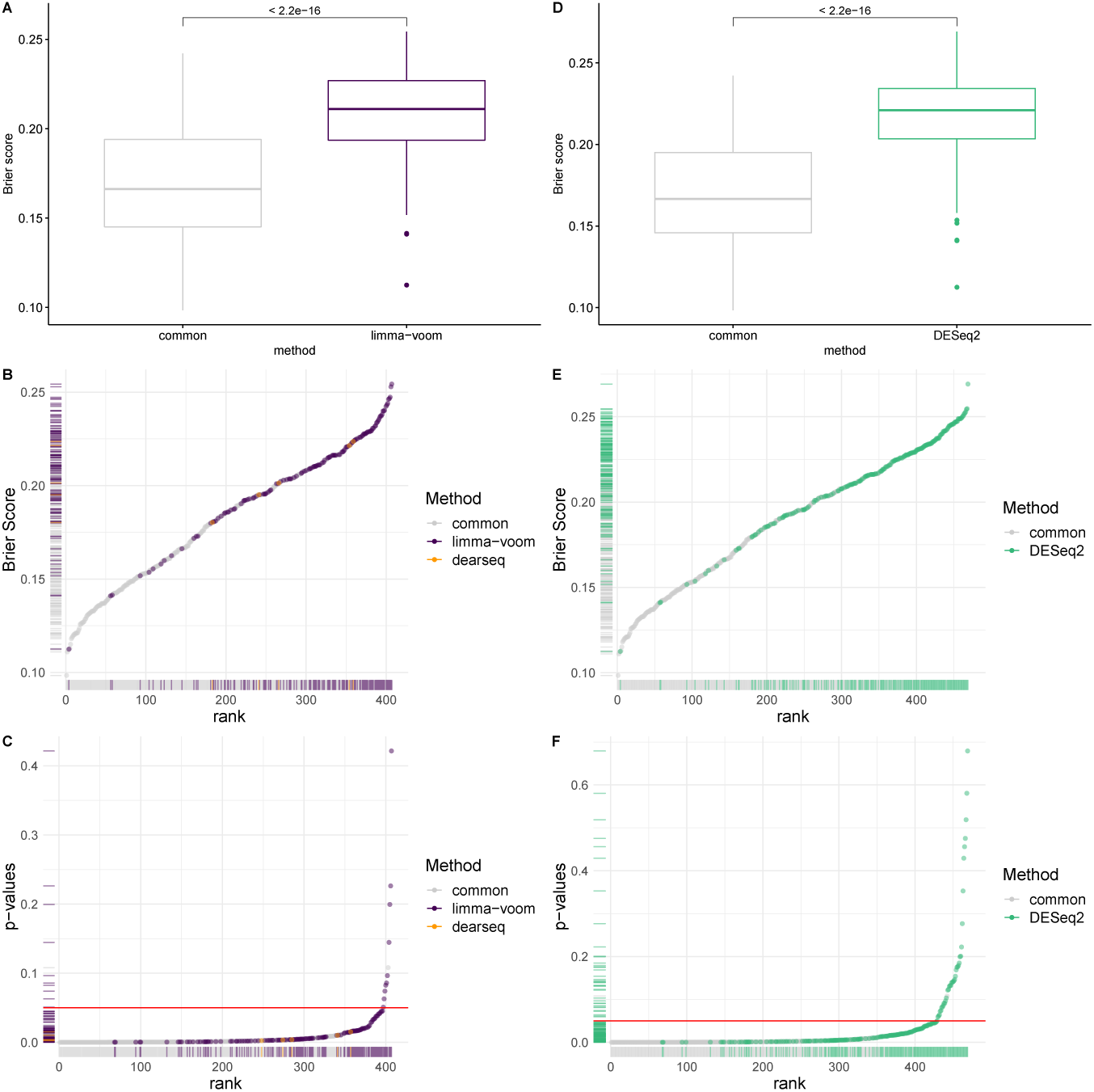
Comparison of the dearseq derived signature to both the DESeq2 and limma-voom derived signatures. a) Boxplots of the Brier scores of the DE genes private to `limma-voom` and the DE genes common to both `dearseq` and `limma-voom`. Note that only 5 genes are identified only by `dearseq` and not `limma-voom`. Therefore we exclude the associated boxplot. b) Univariate Brier scores. The purple points correspond to the DE genes called by `limma-voom` and the grey points to the genes common with `dearseq`. c) Marginal *p*-values. d) Boxplots of the Brier scores of the DE genes private to `dearseq` and the DE genes common to both `dearseq` and `DESeq2`. All genes declared DE by `dearseq` were also declared DE by `DESeq2`. e) Univariate Brier scores. The green points correspond to the DE genes called by `DESeq2` and the grey points to the genes common with `dearseq`. All genes declared DE by `dearseq` were also declared DE by `DESeq2`. f) Marginal *p*-values.

## Discussion

The proposed method `dearseq` represents an innovative and flexible approach for performing gene-wise analysis of RNA-seq gene expression measurements with complex design. As demonstrated in our simulation study, `edgeR`, `DESeq2` or `limma-voom` can all fail to control the Type-I error and the FDR when the sample size increases, while our method behaves correctly. Moreover, the re-analysis of the London Berry Tuberculosis data set revealed that the differentially expressed genes identified by `dearseq` are highly predictive of active Tuberculosis status, while results from the three state-of-the-art methods (including the original `edgeR` analysis) likely include numerous false positives.

It is important to note that `edgeR`, `DESeq2` or `limma-voom` will not systematically have inflated FDR. As illustrated by our simulation studies, there are some scenarios in which, for some given sample sizes, they control the FDR adequately. However, we have shown here that this is no guarantee, and in practice it is very difficult to know under which circumstances a data analysis is taking place.

Because `dearseq` solely relies on the Central Limit Theorem convergence for its asymptotic test to work, it guarantees a control of the FDR without needing any model to hold as long as the sample size is large enough. For lower sample sizes, where convergence is not reached, a robust permutation test can be used instead. Using Phipson & Smyth’s [21] correction, it adequatly controls the FDR regardless of the sample size and exhibits good statistical power in our simulation study (the only trade-off is its increased computation time).

Among the three state-of-art methods compared here, `DESeq2` seems to fail to control the FDR most often. In particular, even under its model assumption of a Negative Binomial distribution for the data, it can suffer from inflated FDR. This seems counter-intuitive as our synthetic data were generated under the Negative Binomial distribution, and this should advantage `DESeq2` and `edgeR`– since both methods assume this model. As has been noted previously, this behavior can be caused by non-uniformity in the distribution of the *p*-values arising from `DESeq2` or `edgeR` (especially when combined with a multiple testing correction such as Benjamini-Hochberg procedure) [22, 12, 23, 24].

DEA can have numerous preprocessing steps, and the various possibilities can complicate the fair comparison of different methods. Since here our primary goal was to compare to the original `edgeR` analysis, we used the `edgeR`-preprocessed data as input to `dearseq`. For `DESeq2` and `limma-voom` we used the raw counts. Indeed both `edgeR` and `DESeq2` assume the input data to be strictly counts (i.e. integers), due to their Negative Binomial distribution assumption, though `edgeR` also has some support for so-called “non-integer counts”. While this seems sensible given the nature of RNA-seq data, recent innovations in RNA-seq alignment methods such as `salmon`[25] or `kallisto`[26] return pseudocounts that are not integers. If the loss of precision is likely not severe when rounding up pseudo-counts, this same limitation prevents the use of already pre-processed (i.e. normalized or transformed data) and forces the DEA practitioner to stick to the specific processing of the methods. In that regard, `dearseq` is extremely flexible and offer to use either raw or transformed data (the default applies a log-cpm transformation similarly to `limma-voom`).

In addition, these methods have been designed to compare two (or multiple) conditions (several treatment regimen), and are not specifically oriented towards grouped or longitudinal data. Therefore there is a need in the broader DEA community for a more flexible method. `dearseq` relies on a general methodology that can easily accommodate more complex designs including gene set analysis while correctly controlling the false discovery rate [27].

## Conclusions

We have demonstrated that the three most popular RNA-seq DEA methods may not guarantee control of the number of false positive in their results, especially when the sample size increases. To exemplify this problematic behavior, we present extensive simulation studies ranging from realistic resampling of real data to synthetic data generation under the models’ assumptions, as well as a re-analysis of a real world data-set. To offer an alternative solution to DEA practitioners, we have developed `dearseq`, a new DEA method that uses a variance component score test to provide a robust, powerful and versatile approach to DEA while avoiding the pitfall of FDR inflation exhibited by the current state-of-the-art methods in certain situations. We also benchmarked this new method alongside the three established methods on both the simulations and the real data analysis to illustrate its excellent performance, both in terms of FDR control and of statistical power.

These results have important implications for the field, as DEA of RNA-seq data has become widespread. The distributional assumptions and model-based inference inherent to `DESeq2`, `edgeR` and `limma-voom` can underestimate the number of false positives in realistic settings. Users should be aware of the possibility of inflated FDR when using these procedures and should consider the use of `dearseq` which gives theoretical and empirical control of the FDR without sacrificing its statistical power. Given the results of both our simulations and our real-world data re-analysis, we thus formulate the following recommendations: i) do not rely on a single DEA method and compare the results across several tools, as this strategy may likely eliminate the majority of false positives; ii) for your main analysis, we recommend using `dearseq` or `limma-voom` over `DESeq2` or `edgeR`. Indeed, `limma-voom` appears to control the FDR adequately as long as your sample size is large enough and the model assumptions (in particular the linearity) are reasonable. On the other hand, `dearseq` ensures an effective control of the FDR regardless of the sample-size (thanks to its permutation test for small sample sizes) and demonstrates good statistical power.

## Methods

The general objective of DEA is to identify genes whose expression is significantly associated with a set of clinically relevant characteristics. `dearseq` is a new DEA framework based on a variance component score test [28, 29, 30], a flexible and powerful test that requires few assumptions to guarantee rigorous control of Type-I and false discovery error rates. The method can be adapted to various experimental designs (comparisons of multiple biological conditions, repeated or longitudinal measurements, integrated supervision by several biomarkers at once). It builds upon recent methodological developments for the analysis of genomic data [31, 32, 30]. Variance component tests offer the speed and simplicity of classical score tests, but potentially gain statistical power by using many fewer degrees of freedom and have been shown to have locally optimal power in some situations [33].

The `dearseq` method comprises 3 steps (with an optional initial normalization):

0. (optional) **normalize** gene expression across samples

1. **Estimate the mean-variance relationship** through a local linear regression borrowing information across all genes

2. **Test** each gene

3. Apply a **multiple-testing correction** controlling the FDR, such as the Benjamini-Hochberg procedure

## Model specification

Let *G* be the total number of observed genes. Let 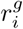 be the raw count of the *g*^*th*^ gene for the *i*^*th*^ sample (*i* = 1,*…, n*). Consider now *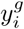* the normalized gene expression (such as log-counts per million, see supplementary materials in additional file 1 for more details). To build a variance component score test statistic, we rely on the following working linear model for each gene *g*:

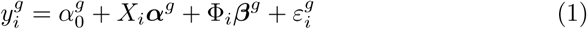

where 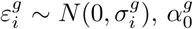 is the intercept, *X*_*i*_ is a vector of covariates to adjust for, and Φ_*i*_ contains the variables for DEA, such as disease status, treatment arm, or other clinical characteristics which are to be associated with gene expression. The parameter of interest is ***β***^*g*^: if ***β***^*g*^ ≠ **0**, then the gene is differentially expressed. The variance of the residuals 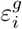 depends on *i* to model the heteroscedasticity inherent to RNA-seq data.

Note that the model presented above is very flexible, and can be easily extended to grouped (e.g. repeated or longitudinal) data to take into account heterogeneity between individuals by adding random effects (see supplementary materials in additional file 1 for more details).

### Estimation of the mean-variance relationship

Because of their count nature, RNA-seq data are intrinsically heteroscedastic. We model this mean-variance relationship through 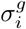. But obviously, this individual variance cannot be estimated from a single observation. Instead, we adopt a strategy similar to `voom` and we gather information across all *G* genes through a local linear regression [34] to estimate 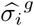 in a rigorous and principled manner (see supplementary materials in additional file 1 for more details).

### Variance component score test statistic estimation

According to the working model (1), a gene is differentially expressed and has its expression associated with the variable(s) of interest in Φ if ***β***^*g*^ ≠ 0. `dearseq` thus tests the following null hypothesis for each gene *g*:

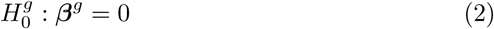

The associated variance component score test statistic can be written as 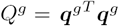 with

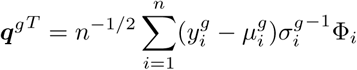

where *μ*_*i*_ is the conditional mean expression given the covariates *X*_*i*_ (see supplementary materials in additional file 1 for more details). Again, this formula can easily generalize to more complex experimental designs such as grouped measurements by incorporating a random-effects covariance matrix (see supplementary materials in additional file 1 for more details).

Because this is a score test, we only need to estimate 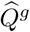 under the null hypothesis of no differential expression. We estimate 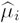 through Ordinary Least Squares. Finally, since a total of *G* tests (with *G* often greater than 10,000) it is absolutely necessary to correct for multiple testing correction, for instance by using the Benjamini-Hochberg procedure.

### Asymptotic and permutation tests

The asymptotic distribution of the test statistic *Q* can be shown to be a mixture of 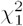 random variables

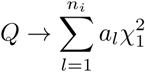

where the mixing coefficients *a*_*l*_ depend on the covariance of ***q*** (see supplementary materials in additional file 1 for details). This asymptotic result rests solely upon the Central Limit Theorem, and this is why `dearseq` is particularly robust to misspecification: the distribution of *Q* is the same whether the model (1) holds or not. Therefore, the Type-I error and the FDR are controlled as long as the Central Limit Theorem is in action (meaning *n* is large enough).

One advantage of using a variance component score test over a regular score test is the gain in statistical power, that comes from exploiting the correlation among ***β***^*g*^ coefficients to potentially reduce the degrees of freedom of the test. Another advantage is its flexibility that can accomodate random effects in the model to test mixed hypotheses (see supplementary materials in additional file 1 for details).

To overcome the shortcomings of this asymptotic test in small samples, we propose to use a permutation test using the same statistic *Q*. Since we are in multiple testing setting it is of the utmost importance to carefully compute the associated *p*-values [21] before applying the Benjamini-Hochberg correction. Finally, in order to preserve statistical power, we use Phipson & Smyth’s correction to account for random permutations (see supplementary materials in additional file 1).

### Availability of data and materials

`dearseq` is freely available at the GitHub repository (https://github.com/borishejblum/dearseq) and in the process of being made available on Bioconductor. The sequence data set from the Singhania *et al.* Tuberculosis study analyzed in this article is accessible from the NCBI GEO database with the primary accession code GSE107991. The code used to analyze the data set and the results are available from the GitHub repository (https://github.com/Mgauth/dearseq_paper).

## Software versions

All computations were run under R v3.5.1 using `DESeq2` package v1.22.1, `edgeR` package v3.24.0, `limma` package v3.38.2, and `dearseq` package v1.0.0.

## Author’s contributions

MG was a major contributor in writing the manuscript and performed the real data re-analysis. DA was a major contributor in writing the manuscript, wrote the original R code for `dearseq`, performed the simulation study under both Negative Binomial and Nonlinear settings and participated in the real data re-analysis. RT was a major contributor in writing the manuscript, participated in the real data analysis and performed the biological interpretation of the results. BH was a major contributor in writing the manuscript, implemented `dearseq` in an R package, performed the resampling simulations and directed the real data re-analysis. All authors read and approved the final manuscript.

### Acknowledgements

MG is supported within the Digital Public Health Graduate’s school, funded by the PIA 3 (Investments for the Future - Project reference: 17-EURE-0019). The project is supported through SWAGR Inria Associate-Team from the Inria@SiliconValley program. Computer time for this study was provided by the computing facilities MCIA (Mésocentre de Calcul Intensif Aquitain) of the Université de Bordeaux and of the Université de Pau et des Pays de l’Adour.

## Additional Files

**Additional file 1 — Supplementary materials**

Supplementary materials as a .PDF describing in details the statistics of `dearseq`, the processing of the data from the London Berry cohort, as well as the simulation settings.

