## Supplementary materials for "dearseq: a variance component score test for RNA-Seq differential analysis that effectively controls the false discovery rate"

### 1 Singhania *et al.* re-analysis

#### 1.1 Input data

RNA-seq data from Singhania *et al.* are publicly available on GEO with the primary accession code GSE107991 for the Berry London cohort. Two files are available:

- raw data : Raw_counts_Berry_London
- `edgeR` preprocessed data : edgeR_normalized_Berry_London

In our re-analysis, we used the `edgeR` preprocessed data to run `limma-voom`, `DESeq2` and `dearseq`. The log fold changes are calculated using the raw data.

**Preprocessing** The matrix of raw counts contains $58,051$ genes and 54 samples. As described in Singhania *et al.*, only genes expressed with counts per million (CPM) $> 2$ in at least five samples were considered and normalized using trimmed mean of M-values (TMM) to remove the library-specific artefacts. The filtering is carried out with `edgeR`. It results in a matrix of normalized counts containing $14,150$ genes and 54 samples (edgeR_normalized_Berry_London).

#### 1.2 Analyses settings

`dearseq` Due to the low sample size for each DEA, the permutation test was used with 1000 permutations. The variable to be tested is the TB group for each of the three comparisons (i.e. TB versus Control, TB versus LTBI and LTBI versus Control). In the absence of covariates, we simply use an intercept.

`DESeq2` We performed the Wald test. The design matrix required was composed of an intercept and the group variable.

`limma-voom` A linear model is fitted to the log2 CPM for each gene. The `voom` step allows to obtain weights for each gene and sample that are passed into `limma`. The design matrix is the same as the two previous methods.

`edgeR` We used the genes signature from Singhania *et al.* supplementary file.

### 2 Detailed simulation settings

#### 2.1 Negative Binomial scenario a)

In this scenario, gene expression is generated from the following Negative Binomial distribution $NB(\mu_{ij}, \tau_{ij})$, such that:

$$E[y_{ij}] = \mu_{ij} \qquad \text{Var}(y_{ij}) = \mu_{ij} + \frac{\mu_{ij}^2}{\tau_{ij}}$$

$$\mu_{ij} = \max\{0, \widetilde{\mu}_{ij}\}$$
$$\widetilde{\mu}_{ij} = \begin{cases} \alpha + b_{i0} + x_i, & j = 1, ..., p_1 \\ \alpha + b_{i0} + x_i + (\beta + b_j) x_i z_i, & j = p_1 + 1, ..., p \end{cases}$$

$$\tau_{ij} \sim \text{exponential}(1), \quad b_j \sim N(0, \sigma_g^2), \quad \alpha = 1000, \quad b_{i0} \sim N(0,1),$$
$$z_i \sim N(0,1), \quad x_i \sim N(\mu_x, 1), \quad \mu_x \sim \text{exponential}(1/10)$$

#### 2.2 Non-linear scenario b)

In this scenario, gene expression is generated according to the following model:

$$\begin{aligned}
y_{ij} &= \min\left\{\max\left\{\frac{\mu_{ij} + \epsilon_{ij}}{10}, 0\right\}, 10^9\right\} \\
\mu_{ij} &= \begin{cases} \eta_{ij} + (\beta + b_j + b_{i1}) z_i, & j = 1, ..., p_1 \\ \eta_{ij}, & j = p_1 + 1, ..., p \end{cases} \\
\eta_{ij} &= \frac{\gamma_{ij} \sum_{i=1}^{n} \gamma_{ij}}{1000n} \\
\gamma_{ij} &= \begin{cases} \nu_{ij} + (\beta + b_j + b_{i1}) z_i, & j = 1, ..., p_1 \\ \nu_{ij}, & j = p_1 + 1, ..., p \end{cases} \\
\nu_{ij} &= \xi_{ij} + \delta_{ij} + x_i \\
\delta_{ij} &\sim N(0, \tau_{ij}^2) \\
\xi_{ij} &= \iota_j \zeta_{ij} + \iota_j \\
\zeta_{ij} &= N(0,1) \\
\iota_j &\sim \text{exponential}(0.01) \\
b_j &\sim N(0, \sigma_g^2) \\
\log(\epsilon_{ij}) &\sim N(0, \sigma_{ij}^2) \\
\sigma_{ij} &\sim \text{exponential}(1)
\end{aligned}$$

and $\tau_j$ is 0.01 times the standard deviation of $\{\xi_{1j}, ..., \xi_{nj}\}$

#### 2.3 SEQC resampling scenario c)

In this scenario, gene expression was generated by randomly sampling among the five "A" replicate samples from the SEQC data. In practice, we used the data provided as supplementary data by Rapaport *et al.* [1]. For a given sample size (4, 8, 16, 20, 50, 100, 150, 200 or 300), each simulated sample was first drawn from the original five real ones and arbitrarily assigned to one of the two mock comparison groups. Then, random noise was added (using a multivariate Gaussian distribution centered on 0 with a covariance matrix for all the genes estimated from the five real original samples) in order to obtain different values for each simulated samples. Finally, values were rounded to the nearest integer and truncated at 0 (included), in order to emulate count data. Since the five A samples are replicates, such simulated samples were homogeneous and did not feature any truly DE gene.

### 3 dearseq

This section details the statistical grounds of `dearseq`. We present `dearseq`in its most general form for the benefit of users of the method and software who may have more complex data. This most general form is given in Sections 3.2, 3.3, 3.4, and 3.5. Simplifications that connect the general development with the specific analysis in the main text are given in 3.6.

#### 3.1 Normalized gene expression

The `dearseq` methodology assumes that the gene expression measurement are comparable across samples. As this is not always the case with raw RNA-seq counts (due to technical effects for instance), a normalization step is often required. `dearseq` does not assume any specific normalization and can work with any kind of quantitative variables.

#### 3.2 Most general modeling framework for `dearseq`

In this section, we demonstrate how `dearseq`can be used to analyze longitudinal, grouped, or repeated measurements. Simplifications for a single observation per individual are given in 3.6. Let $P$ be the total number of observed genes. Let $y_{ij}^g$ be the normalized gene expression of the $g^{th}$ gene for the $i^{th}$ sample at the $j^{th}$ measure, for $i = 1, \dots, n$, $j = 1, \dots, n_i$. Further let $\boldsymbol{\phi}_{ij}$ be the quantity we are interested in testing. In the main analysis in the text, $\boldsymbol{\phi}_{ij}$ would correspond to TB status (active TB, LTBI, or control). In other cases, it might include treatment arm in a clinical trial, a quantitative measure of disease, or any combination of continuous or binary measures that are under study.

To build a variance component score test statistic, we rely on the following working model for each gene $g$:

$$y_{ij}^g = \alpha_0^g + X_{ij}^T \boldsymbol{\alpha}^g + \boldsymbol{\phi}_{ij}^T \boldsymbol{\beta}^g + \boldsymbol{\phi}_{ij}^T \boldsymbol{\xi}_i^g + \epsilon_{ij}^g, \tag{1}$$

which can be factorized into matrix form as:

$$\boldsymbol{y}_i^g = \boldsymbol{\alpha}_0^g + X_i \boldsymbol{\alpha}^g + \Phi_i \boldsymbol{\beta}^g + \Phi_i \boldsymbol{\xi}_i^g + \boldsymbol{\epsilon}_i^g, \tag{2}$$

where, $\boldsymbol{y}_i = (y_{i1}, \dots, y_{in_i})$ is a $n_i \times 1$ vector of normalized gene expression measurements, $\boldsymbol{\epsilon}_i \sim N(0, \Sigma_i)$ is a $n_i \times 1$ vector of measurement error, $\boldsymbol{\alpha}_0$ is a

$n_i \times 1$ vector of intercepts, $\boldsymbol{\alpha}$ is a $p \times 1$ vector of fixed effects, $\boldsymbol{\beta}$ and $\boldsymbol{\xi}_i \sim N(0, \Sigma_\xi)$ are respectively a $m \times 1$ vector of fixed effects and a $m \times 1$ vector of individual-level random effects of the variables of interest, $X_i$ and $\Phi_i$ are the associated $n_i \times p$ matrix of covariates and $n_i \times m$ matrix of the variables of interest. $\Sigma_\xi$ is the $m \times m$ covariance matrix of $\boldsymbol{\xi}_i$. $\Sigma_i$ is the $n_i \times n_i$ covariance matrix of measurement errors. $\boldsymbol{\xi}_i$ and $\boldsymbol{\epsilon}_i$ are assumed to be independent. Note that, to take into account the correlation between the different measurements of the same individual we use a random effect in the model. In addition, it is important to note that the variance of the residuals depends on $i$ and $j$ to model the heteroscedasticity of the data. This means that each measure of each individual has a different variance.

#### 3.3 Estimating the mean-variance relationship

A key step in our method is the estimation of $\Sigma_i = \text{diag}(\sigma_{i1}^2, \ldots, \sigma_{in_i}^2)$ $\forall i$ and for each gene, which will be useful for calculating the test statistic. Because of the intrinsic heteroscedasticity of the data, the variance of the residuals depends on $i$ and $j$, so, the variance cannot be estimated only from a single observation, especially with conventional estimators. To approximate the mean-variance relationship in $\mathbf{y}^g$, we use information from all P genes. Let $v_{ij}^g = Var(y_{ij}^g \mid X_{ij}, \boldsymbol{\xi}_i^g)$ and $m_{ij}^g = E(y_{ij}^g \mid X_{ij}, \boldsymbol{\xi}_i^g)$ respectively the variance and the mean of gene $g$ for sample $i$ and measure $j$ given the covariates and the random effects. We assume that $v_{ij}^g$ may be modeled as a function of its mean $m_{ij}^g$. To save computational time and reduce the number of points used in the nonparametric fit, one could follow Law *et al.* [2] and model the mean-variance relationship at the gene level. Specifically, $v^g = \omega(m^g) + e^g$ for some unknown function $\omega(.)$ and errors which follow the moment conditions $E(e^g) = 0, V(e^g) = \tau^2$, $\tau > 0$. Thus, we used a local linear regression proposed by Wasserman [3] which offers good asymptotic convergence:

$$\tilde{S}_{nd}(x) = \sum_g K\left(\frac{m^g - x}{h}\right)(m^g - x)^d, \text{ for } d = 1, 2$$

$$\tilde{b}^g(x) = K\left(\frac{m^g - x}{h}\right)(\tilde{S}_{n2}(x) - (m^g - x)\tilde{S}_{n1}(x)),$$

$$\tilde{l}^g(x) = \frac{\tilde{b}^g(x)}{\sum_g \tilde{b}^g(x)}, \quad \tilde{\omega}_n(x) = \sum_g \tilde{l}^g(x) v^g \tag{3}$$

for some kernel function $K(.)$ and bandwidth $h > 0$. Standard cross-validation techniques may be used to select $h$ in practice.

Because the mixed effects model (1) may be computationally costly, to obtain the mean-variance relationship, we further simplify to a fixed effects model,

$$\boldsymbol{y}_i^g = \alpha_0^g + X_i^T \boldsymbol{\alpha}^g + \Phi_i^T \boldsymbol{\beta}^g + \tilde{\varepsilon}_i^g. \tag{4}$$

Based on this model, the mean-variance relationship could be estimated by $\widehat{\omega}_n(x) = \tilde{\omega}_n(x)_{|m^g=\widehat{m}^g, v^g=\widehat{v}^g}$ with the estimate of the mean $\widehat{m}^g = n^{-1}\sum_{i=1}^{n} n_i^{-1}\sum_{j=1}^{n_i}\widehat{\alpha}_0^g + X_{ij}^T\widehat{\boldsymbol{\alpha}}^g + \Phi_{ij}^T\widehat{\boldsymbol{\beta}}^g$ and the estimate of the variance $\widehat{v}^g = n^{-1}\sum_{i=1}^{n} n_i^{-1}\sum_{j=1}^{n_i}(y_{ij}^g - \widehat{\alpha}_0^g - X_{ij}^T\widehat{\boldsymbol{\alpha}}^g - \Phi_{ij}^T\widehat{\boldsymbol{\beta}}^g)^2$ where " $\widehat{}$ " is the Ordinary Least Square Estimator.

Now that we have the estimate of $\omega_n$, we can calculate the variance estimate of $y_{ij}^g$ for all P genes as: $(\widehat{\sigma}_{ij}^g)^2 = \widehat{\omega}_n(\widehat{m}_{ij}^g)$ with $\widehat{m}_{ij}^g = \widehat{\alpha}_0^g + X_{ij}^T\widehat{\boldsymbol{\alpha}}^g + \Phi_{ij}^T\widehat{\boldsymbol{\beta}}^g$.

##### 3.4 Test statistic

In this section, we derive a variance component score test statistic for the effects of interest. For the sake of simplicity, we omit the gene index $g$ in the following, bear in mind that a test is carried out for each gene $g$.

According to the model (2), the null hypothesis of no effect of interest is:

$$H_0 : \text{“}\boldsymbol{\beta} = 0 \text{ and } \Sigma_\xi = 0\text{”} \tag{5}$$

If the variance-covariance matrix of the random effects is identically zero then the random effects $\boldsymbol{\xi}_i$ are also identically zero for all $i$. If at the same time, $\boldsymbol{\beta} = 0$ then the expression of the gene will not be significantly associated with the variables of interest $\Phi_i$.

Under the working model (2), for all $i$, we will distinguish the effects of covariates and the effects of variables of interest on gene expression by posing $\boldsymbol{\mu}_i = \boldsymbol{\alpha}_0 + X_i\boldsymbol{\alpha}$ and $\boldsymbol{\theta}_i = \boldsymbol{\beta} + \boldsymbol{\xi}_i$. We write $\boldsymbol{\theta}_i = \eta\boldsymbol{\nu}_i = \eta(\boldsymbol{\gamma} + \boldsymbol{\zeta}_i)$ with $\boldsymbol{\nu}_i \sim N(0, \Sigma_\nu), \boldsymbol{\gamma} \sim N(0, I), \boldsymbol{\zeta}_i \sim N(0, \Sigma_\zeta), \Sigma_\nu = I + \Sigma_\zeta$. $\boldsymbol{\nu}_i$ is the nuisance parameter. We can rewrite the null hypothesis as $H_0$ : "$\eta = 0$" and the model as $\boldsymbol{y}_{\mu_i} = \eta\Phi_i\boldsymbol{\nu}_i + \boldsymbol{\varepsilon}_i$ with $\boldsymbol{y}_{\mu_i} = \boldsymbol{y}_i - \boldsymbol{\mu}_i$ the centered outcome. Then, $\boldsymbol{y}_{\mu_i}|\boldsymbol{\nu}_i \sim N(\eta\Phi_i\boldsymbol{\nu}_i, \Sigma_i)$. We write the likelihood of $\boldsymbol{y}_{\mu_1}, \ldots, \boldsymbol{y}_{\mu_n}|\boldsymbol{\nu}_i$:

$$\begin{aligned}\mathcal{L}(\eta) &= \mathcal{L}(\boldsymbol{y}_{\mu_1}, \ldots, \boldsymbol{y}_{\mu_n}, \eta|\boldsymbol{\nu}_i)\\ &= \prod_{i=1}^{n}\frac{1}{(2\pi)^{n_i/2} \mid \Sigma_i \mid^{1/2}}\\ &\quad exp\left(-\frac{1}{2}(\boldsymbol{y}_{\mu_i} - \eta\Phi_i\boldsymbol{\nu}_i)^T\Sigma_i^{-1}(\boldsymbol{y}_{\mu_i} - \eta\Phi_i\boldsymbol{\nu}_i)\right)\end{aligned}$$

Then, we derive a variance component score test. It has the advantage of avoiding the estimation of $\boldsymbol{\beta}$ and $\boldsymbol{\xi}_i$ because it only requires estimating the model under the null.

We write the likelihood of $\boldsymbol{y}_{\mu_1}, ..., \boldsymbol{y}_{\mu_n}|\boldsymbol{\nu}_i$:

$$\begin{aligned}\mathcal{L}(\eta) &= \mathcal{L}(\boldsymbol{y}_{\mu_1}, ..., \boldsymbol{y}_{\mu_n}, \eta|\boldsymbol{\nu}_i) \\ &= \prod_{i=1}^{n} \frac{1}{(2\pi)^{n_i/2} \mid \Sigma_i \mid^{1/2}} exp\left(-\frac{1}{2}(\boldsymbol{y}_{\mu_i} - \eta\Phi_i\boldsymbol{\nu}_i)^T \Sigma_i^{-1} (\boldsymbol{y}_{\mu_i} - \eta\Phi_i\boldsymbol{\nu}_i)\right)\end{aligned}$$

The score being null, we follow the argument of Commenges and Andersen [4] by considering $\lim_{\eta\to 0} \frac{\partial}{\partial(\eta^2)} log(\mathcal{L}^*(\eta))$ to obtain the expression of the test statistic. Let $\mathcal{L}^*(\eta)$ be the likelihood of $\boldsymbol{y}_{\mu_1}, ..., \boldsymbol{y}_{\mu_n}$ such as:

$$\mathcal{L}^*(\eta) = \mathcal{L}(\boldsymbol{y}_{\mu_1}, ..., \boldsymbol{y}_{\mu_n}; \eta) = \mathbb{E}[\mathcal{L}(\boldsymbol{y}_{\mu_1}, ..., \boldsymbol{y}_{\mu_n}; \eta|\boldsymbol{\nu}_i) \mid \mathbb{V}],$$

with $\mathbb{V} = \{\boldsymbol{V}_i = (\boldsymbol{y}_{\mu_i}^T, X_i^T, \Phi_i^T)^T\}_{i=1}^n$
Then,

$$\begin{aligned}\lim_{\eta\to 0} \frac{\partial log(\mathcal{L}^*(\eta))}{\partial(\eta^2)} &= \lim_{\eta\to 0} \frac{1}{2\eta\mathcal{L}^*(\eta)} \frac{\partial\mathcal{L}^*(\eta)}{\partial\eta} \\ &= \lim_{\eta\to 0} \frac{1}{2\mathcal{L}^*(\eta)} \left[\eta^{-1}\frac{\partial\mathcal{L}^*(0)}{\partial\eta} + \frac{\partial^2\mathcal{L}^*(0)}{\partial\eta^2} + o(1)\right] \\ &= \frac{1}{2\mathcal{L}^*(0)} \left[\frac{\partial^2\mathcal{L}^*(0)}{\partial\eta^2} + o(1)\right]\end{aligned}$$

Thus, removing 1/2 for the sake of simplicity, we have :

$$\begin{aligned}\mathcal{L}^*(0)^{-1}\frac{\partial^2\mathcal{L}^*(0)}{\partial\eta^2} + o(1) &= \mathcal{L}^*(0)^{-1}\frac{\partial}{\partial\eta}\left(\frac{\partial\mathcal{L}^*(0)}{\partial\eta}\right) + o(1) \\ &= \frac{\partial^2 log\mathcal{L}^*(0)}{\partial\eta^2} + o(1) \\ &= \mathbb{E}\left[\frac{\partial^2 log\mathcal{L}(0)}{\partial\eta^2}|\mathbb{V}\right] + o(1) \\ &= \mathbb{E}\left[\frac{\partial}{\partial\eta}\left(\frac{\partial log\mathcal{L}(0)}{\partial\eta}\right)|\mathbb{V}\right] + o(1) \\ &= \mathbb{E}\left[\frac{\frac{\partial^2\mathcal{L}(0)}{\partial\eta^2}\mathcal{L}(0) - \frac{\partial\mathcal{L}(0)}{\partial\eta}\frac{\partial\mathcal{L}(0)}{\partial\eta}}{\mathcal{L}(0)^2}|\mathbb{V}\right] + o(1) \\ &= \mathbb{E}\left[\frac{\frac{\partial^2\mathcal{L}(0)}{\partial\eta^2}}{\mathcal{L}(0)} - \left(\frac{\frac{\partial\mathcal{L}(0)}{\partial\eta}}{\mathcal{L}(0)}\right)^2 |\mathbb{V}\right] + o(1) \\ &= \mathbb{E}\left[\frac{\partial^2 log\mathcal{L}(0)}{\partial\eta^2} - \left(\frac{\partial log\mathcal{L}(0)}{\partial\eta}\right)^2 |\mathbb{V}\right] + o(1) \\ &= \mathbb{E}\left[\frac{\partial^2 log\mathcal{L}(0)}{\partial\eta^2}|\mathbb{V}\right] + \mathbb{E}\left[\left(\frac{\partial log\mathcal{L}(0)}{\partial\eta}\right)^2 |\mathbb{V}\right] + o(1)\end{aligned}$$

Then standardizing by $n$,

$$\lim_{\eta\to 0} n^{-1}\frac{\partial log(\mathcal{L}^*(\eta))}{\partial(\eta^2)} = n^{-1}\mathbb{E}\left[\left(\frac{\partial log\mathcal{L}(0)}{\partial\eta}\right)^2|\mathbb{V}\right] + \text{constant} + o(1)$$

because

$$n^{-1}\frac{\partial^2 log(\mathcal{L}(\eta))}{\partial\eta^2} = n^{-1}\sum_{i=1}^{n} - (\Phi_i\boldsymbol{\nu}_i)^T\Sigma_i^{-1}\Phi_i\boldsymbol{\nu_i} = \text{constant}$$

Yet,

$$\begin{aligned}
\frac{\partial log(\mathcal{L}(\eta))}{\partial\eta} &= \sum_{i=1}^{n}\frac{\partial}{\partial\eta} - \frac{n_i}{2}log(2\pi) - \frac{1}{2}log(\mid\Sigma_i\mid) - \frac{1}{2}(\boldsymbol{y}_{\mu_i} - \eta\Phi_i\boldsymbol{\nu}_i)^T\Sigma_i^{-1}(\boldsymbol{y}_{\mu_i} - \eta\Phi_i\boldsymbol{\nu}_i) \\
&= \sum_{i=1}^{n}\frac{\partial}{\partial\eta} - \frac{n_i}{2}log(2\pi) - \frac{1}{2}log(\mid\Sigma_i\mid) - \frac{1}{2}(\boldsymbol{y}_{\mu_i}^T\Sigma_i^{-1}\boldsymbol{y}_{\mu_i} - \boldsymbol{y}_{\mu_i}^T\Sigma_i^{-1}\eta\Phi_i\boldsymbol{\nu}_i \\
&- \eta(\Phi_i\boldsymbol{\nu}_i)^T\Sigma_i^{-1}\boldsymbol{y}_{\mu_i} + \eta^2(\Phi_i\boldsymbol{\nu}_i)^T\Sigma_i^{-1}\Phi_i\boldsymbol{\nu}_i) \\
&= \sum_{i=1}^{n} - \frac{1}{2}\left(-\boldsymbol{y}_{\mu_i}^T\Sigma_i^{-1}\Phi_i\boldsymbol{\nu}_i - (\Phi_i\boldsymbol{\nu}_i)^T\Sigma_i^{-1}\boldsymbol{y}_{\mu_i} + 2\eta(\Phi_i\boldsymbol{\nu}_i)^T\Sigma_i^{-1}\Phi_i\boldsymbol{\nu}_i\right)
\end{aligned}$$

and

$$\begin{aligned}
\frac{\partial log(\mathcal{L}(\eta))}{\partial\eta}\mid_{\eta=0} &= \sum_{i=1}^{n} - \frac{1}{2}(-\boldsymbol{y}_{\mu_i}^T\Sigma_i^{-1}\Phi_i\boldsymbol{\nu}_i - (\Phi_i\boldsymbol{\nu}_i)^T\Sigma_i^{-1}\boldsymbol{y}_{\mu_i}) \\
&= \sum_{i=1}^{n}\frac{1}{2}(\boldsymbol{y}_{\mu_i}^T\Sigma_i^{-1}\Phi_i\boldsymbol{\nu}_i + \boldsymbol{y}_{\mu_i}^T\Sigma_i^{-1}\Phi_i\boldsymbol{\nu}_i) \\
&= \sum_{i=1}^{n}\boldsymbol{y}_{\mu_i}^T\Sigma_i^{-1}\Phi_i\boldsymbol{\nu}_i
\end{aligned}$$

So,

$$
\begin{aligned}
n^{-1}\mathbb{E}\left[\left(\frac{\partial log\mathcal{L}(0)}{\partial\eta}\right)^2|\mathbb{V}\right] &= n^{-1}Var\left[\frac{\partial log\mathcal{L}(0)}{\partial\eta} \mid \mathbb{V}\right] \\
&= n^{-1}Var\left[\sum_{i=1}^{n}\boldsymbol{y}_{\mu_i}^T\Sigma_i^{-1}\Phi_i\boldsymbol{\nu}_i \mid \mathbb{V}\right] \\
&= n^{-1}Var\left[\sum_{i=1}^{n}\boldsymbol{y}_{\mu_i}^T\Sigma_i^{-1}\Phi_i(\boldsymbol{\gamma}+\boldsymbol{\zeta}_i) \mid \mathbb{V}\right] \\
&= n^{-1}Var\left[\left(\sum_{i=1}^{n}\boldsymbol{y}_{\mu_i}^T\Sigma_i^{-1}\Phi_i\right)\boldsymbol{\gamma} \mid \mathbb{V}\right] + n^{-1}Var\left[\sum_{i=1}^{n}\boldsymbol{y}_{\mu_i}^T\Sigma_i^{-1}\Phi_i\boldsymbol{\zeta}_i \mid \mathbb{V}\right] \\
&= n^{-1}\left(\sum_{i=1}^{n}\boldsymbol{y}_{\mu_i}^T\Sigma_i^{-1}\Phi_i\right)\left(\sum_{i=1}^{n}\boldsymbol{y}_{\mu_i}^T\Sigma_i^{-1}\Phi_i\right)^T + n^{-1}\sum_{i=1}^{n}\boldsymbol{y}_{\mu_i}^T\Sigma_i^{-1}\Phi_i\Sigma_\zeta\Phi_i^T\Sigma_i^{-1}\boldsymbol{y}_{\mu_i} \\
&= \left(n^{-1/2}\sum_{i=1}^{n}\boldsymbol{y}_{\mu_i}^T\Sigma_i^{-1}\Phi_i\right)\left(n^{-1/2}\sum_{i=1}^{n}\boldsymbol{y}_{\mu_i}^T\Sigma_i^{-1}\Phi_i\right)^T + \text{constant} + o(1) \\
&= \boldsymbol{q}^T\boldsymbol{q}
\end{aligned}
$$

Let $Q$ be the variance component score test statistic such as $Q = \boldsymbol{q}^T\boldsymbol{q}$ with

$$
\boldsymbol{q}^T = n^{-1/2}\sum_{i=1}^{n}\boldsymbol{y}_{\mu_i}^T\Sigma_i^{-1}\Phi_i = n^{-1/2}\sum_{i=1}^{n}(\boldsymbol{y}_i - \mu_i)^T\Sigma_i^{-1}\Phi_i
$$

Considering that we have a consistent estimator of $\Sigma_i$, we still must provide estimates of $\alpha_0$ and $\boldsymbol{\alpha}$. A natural way to estimate these quantities, given the heteroscedasticity in $\boldsymbol{y}$, is to fit a weighted mixed effects model. The weights are taken to be $\boldsymbol{w}_i = \text{diag}(\widehat{\Sigma}_i)^{-1}$. However, to avoid excessive computation time, instead of estimating the full mixed effects model from (2), we may fit a simpler fixed effects model (4), from which we can obtain estimates of $\alpha_0$ and $\boldsymbol{\alpha}$.

##### 3.5 Test statistic limiting distribution

We have $Q = \boldsymbol{q}^T\boldsymbol{q}$. We note $\Gamma = Cov(\boldsymbol{q})$. Then, we can write:

$$
Q = \boldsymbol{q}^T\Gamma^{-1/2}\Gamma\Gamma^{-1/2}\boldsymbol{q}
$$

The matrix $\Gamma$ being square and diagonal, we carry out a singular value decomposition of $\Gamma$:

$$
Q = \boldsymbol{q}^T\Gamma^{-1/2}UAU^T\Gamma^{-1/2}\boldsymbol{q},
$$

where $U$ is an orthogonal matrix of eigen vectors of $\Gamma$, $A$ is a diagonal matrix of eigen values of $\Gamma$.

We take $u = \Gamma^{-1/2}\boldsymbol{q}$, with $\boldsymbol{q}^T = n^{-1/2}\sum_{i=1}^{n}(\boldsymbol{y}_i - \boldsymbol{\mu}_i)^T\Sigma_i^{-1}\Phi_i\Sigma_\nu^{1/2}$, which give us:

$Q = u^T U A U^T u$. Under the normal residual hypothesis, $u$ immediately follows a standard normal distribution, in which case it is not necessary to apply the central limit theorem. Nevertheless, the variance component test is intended to be robust to the mispecification of the model, i.e., if the normal residual assumption is not verified. Therefore, we propose an asymptotic test to ensure its robustness against any data distribution, for example a negative binomial. This is one of the reasons why we propose the use of permutations when the number of individuals is considered too small or simply to ensure the reliability of the test. Then,

$$\begin{aligned}
\mathbb{E}(u) = \Gamma^{-1/2}\mathbb{E}(\boldsymbol{q}^T) &= \Gamma^{-1/2}\mathbb{E}(n^{-1/2}\sum_{i=1}^{n}(\boldsymbol{y}_i - \boldsymbol{\mu}_i)^T \Sigma_i^{-1}\Phi_i \Sigma_\nu^{1/2}) \\
&= \Gamma^{-1/2} n^{-1/2}\sum_{i=1}^{n} \underbrace{\mathbb{E}(\boldsymbol{y}_i^T - \boldsymbol{\mu}_i^T)}_{=0} \Sigma_i^{-1}\Phi_i \\
&= 0
\end{aligned}$$

and

$$\begin{aligned}
Cov(u) &= Cov(\Gamma^{-1/2} q^T) \\
&= (\Gamma^{-1/2})^T Cov(q) \Gamma^{-1/2} \\
&= \Gamma^{-1/2}\Gamma\Gamma^{-1/2} \\
&= \Gamma^{-1/2}\Gamma^{1/2} I_{n_i} \Gamma^{1/2}\Gamma^{-1/2} \\
&= I_{n_i}
\end{aligned}$$

By the central limit theorem, $u$ asymptotically follows a multivariate standard normal distribution. $U$ being orthonormal, $U^T u$ also asymptotically follows a multivariate standard normal distribution. So, $u^T U A U^T u = \sum_{k=1}^{n_i} a_k (u_k^*)^2$ where $u_k^*$ is an element of the asymptotic multivariate standard normal distribution of $U^T u$ and $a_k$ is an eigen value of $\Gamma$. So, it follows that $Q \underset{+\infty}{\sim} \sum_{k=1}^{n_i} a_k \chi_1^2$. Let $\widehat{Q}$ be the estimate of $Q$. Because $Q$ and $\widehat{Q}$ are asymptotically equivalent, $\widehat{Q} \underset{+\infty}{\sim} \sum_{k=1}^{n_i} \widehat{a}_k \chi_1^2$. (See Agniel and Hejblum (2017) for the proof.)

#### 3.6 Simplification when the measurements are not repeated

In this section, we detail how the generic formulation of the variance component score test simplifies into the form given in the main manuscript when the data are not repeated. When there is only one observation per individual and only one variable of interest (i.e., $\phi_{ij}$ is a scalar), the variance component score test simplifies to a standard score test. When there are multiple variables of interest and $\Phi_{ij}$ is a vector, the variance component score test may gain additional statistical power thanks to its exploitation of potential correlation among the tested variables (through its chi-square mixture asymptotics - see section 3.5 for more details). Here, we assume that the data are not grouped (e.g. repeated or longitudinal) and therefore the index $j$ has to be removed, as used in the main manuscript. Thus, let $y_i^g$ be the normalized gene expression of the $g^{th}$ gene for the $i^{th}$ sample. The working model is written as follows:

$$y_i^g = \alpha_0^g + X_i \boldsymbol{\alpha}^g + \Phi_i \boldsymbol{\beta}^g + \varepsilon_i^g \tag{6}$$

where $\varepsilon_i^g \sim N(0, (\sigma_i^g)^2)$, $\alpha_0^g$ is the intercept, $X_i$ is a vector of $p$ observations from covariates that needs to be adjusted, $\boldsymbol{\alpha}^g$ is the corresponding vector of $p$ fixed effects, $\Phi_i$ is a vector of $m$ observations from the variables of interest (with whom the expression association is tested), and $\boldsymbol{\beta}^g$ is the corresponding $m$ vector of fixed effects associated to those variables of interest. The variance of the residuals depends on $i$ to model the heteroscedasticity of the observations $\boldsymbol{y}$.

According to the working model (2), a gene has its expression associated with the variable(s) of interest in $\Phi$ if $\boldsymbol{\beta}^g \neq 0$. `dearseq` thus tests the following null hypothesis:

$$H_0^g : \boldsymbol{\beta}^g = 0$$

The associated variance component score test statistic can be written as $Q^g = \boldsymbol{q}^{gT} \boldsymbol{q}^g$ with

$$\boldsymbol{q}^{gT} = n^{-1/2} \sum_{i=1}^{n} (y_i^g - \mu_i^g)(\sigma_i^g)^{-1} \Phi_i,$$

where $\mu_i$ is the conditional mean normalized expression given the covariates $X_i$.

#### 3.7 Asymptotic and permutation tests

When $n$ is sufficiently large, we propose an asymptotic test. The asymptotic distribution of the test statistic $Q$ is a mixture of $\chi_1^2$ random variables, i.e. $Q \to \sum_{l=1}^{n_i} a_l \chi_1^2$ where the mixing coefficients $a_l$ depend on the covariance of $\boldsymbol{q}$. When $n$ is very small, relying on the limiting distribution may not be adequate. To overcome this difficulty, we provide a permutation alternative to our asymptotic test. Permutations can be used to estimate the empirical distribution of $\widehat{Q}$ under the null hypothesis. Indeed, permutation tests are attractive because the only assumption we make is that the observations are independent and identically distributed under the null. As explained Phipson and Smyth [5], it's essential to notice that permutation $p$-values that are really estimates of $p$-values, i.e., $\widehat{p}$-values, can lead to $\widehat{p}$-values exactly equal to zero. However, it is senseless to obtain $\widehat{p}$-values equal to zero when all permutations were enumerated, therefore it is not accurate to assume that the $\widehat{p}$-value can be reduced to zero by taking a smaller subset of all the permutations. So, estimating the $p$-value by $B/m$ where $B$ is the number of permutations for which the associated test statistics are at least as extreme as the observed one can be misleading.

Considering our model, the observations of a given individual $i$ are exchangeable under the null, regardless of sampling measure. We assume that an independent random sample of $m$ permutations is drawn with replacement such as

$\boldsymbol{y}_i^* \in \mathbb{R}^{n_i}, y_{ij}^* = y_{i\sigma(j)}$ with $\sigma \in Perm\{1, \ldots, n_i\}$. We generate $m$ test statistics which can contain repeat values, including the original observed value $t_{obs}$. Let $B$ be the number of permutations for which the $m$ test statistics are at least as extreme as $t_{obs}$, $m_t$ be all possible distinct permutations, $B_t$ be the unknown total number of possible distinct test statistics exceeding $t_{obs}$, and $p_t = (B_t + 1)/(m_t + 1)$ be the ideal permutation $p$-value which is obviously unknown. If the null hypothesis is true, then $B_t$ follows a discrete uniform distribution on the integers $0, \ldots, m_t$ . Conditional on $B_t = b_t$ , $B$ follows a binomial distribution $\mathcal{B}(m, p_t)$. An approximation of this quantity can be calculated by:

$$p_e = \frac{b+1}{m+1} - \int_0^{0.5/m_t+1} F(b; m, p_t), \tag{7}$$

$F$ is the cumulative probability function of the binomial distribution.
